# Unilateral naris occlusion with custom-made plugs for reversible olfactory deprivation in juvenile and adult mice

**DOI:** 10.64898/2026.11.03.763032

**Authors:** Ana Dorrego-Rivas, Sonu Peedikayil-Kurien, Li Huang, Tess C. Stanley, Elisa Galliano

## Abstract

Deprivation paradigms that reduce environmental input have been successfully applied across multiple species and sensory modalities, including vision, audition, somatosensation, and olfaction in order to understand how neurons and circuits respond to loss of sensory input. One notable advantage of studying sensory deprivation in the rodent olfactory system is the ease and control with which smell loss can be induced. Various methods of olfactory deprivation have long been available and rely on blocking a nostril, either through invasive and often non-reversible approaches such as chemical ablation or surgical procedures, or through reversible approaches such as unilateral naris occlusion using custom-made nose plugs. Here, we provide a systematic description of this well-established method, including its construction from commercially available materials, procedures, validation, and reliability in post-weaning mice, thereby providing a detailed and reproducible framework for controlled, reversible olfactory deprivation.

## INTRODUCTION

Neuroscience has long investigated the effects of changes in sensory inputs on brain structure and function by depriving a sense through surgical, chemical, or mechanical interventions targeting the peripheral organ, either during development or adulthood (Hubel and Wiesel, 1963; Coppola, 2012; Huang et al., 2024). Compared with sensory enrichment, where heterogeneous stimuli can differentially engage specific circuits and complicate interpretation due to sampling biases unless cells responding to a known stimulus are genetically labelled (Horgue et al., 2022), deprivation provides a uniform manipulation of sensory drive. Among sensory modalities, olfaction is pivotal for most mammals, including humans, with its loss having profound impacts on physical and mental health, as highlighted by the COVID-19 pandemic (Kohli et al., 2016; Boesveldt et al., 2017; Xydakis et al., 2021).

Mice, one of the most widely used animal models, rely heavily on olfaction to shape their behaviour and prevention of odour sampling through their nostrils provides a powerful approach to understanding how sensory activity sculpts the structure and function of downstream central circuits (Saghatelyan et al., 2005; George et al., 2022; Coppola and Reisert, 2023).

Traditionally, olfactory deprivation has been achieved surgically, chemically, or mechanically using nose plugs. The specific approach determines whether deprivation can be performed bilaterally, eliminating sensory input from both nostrils and both olfactory bulbs (OBs), or unilaterally, affecting only one side. The OB is the first central relay in the olfactory pathway, where olfactory sensory neuron (OSN) axons from the olfactory epithelium (OE) in the nose synapse within glomeruli onto the dendrites of local interneurons and of mitral and tufted cells, which project to the olfactory cortex and higher processing areas. Because the olfactory system is largely ipsilateral during these early stages with each nostril projecting to the ipsilateral OB (Grimaud et al., 2024), unilateral deprivation selectively reduces sensory input to a single OB. To achieve full loss of olfactory input to both olfactory bulbs, approaches include surgical interventions that axotomise OSNs (Mandairon et al., 2003), or chemical approaches that damage or ablate OSNs using systemically administered drugs such as methimazole (Kikuta et al., 2015; Browne et al., 2022) or via nasal lavages of agents such as zinc sulphate (McBride et al., 2003). However, these approaches produce only transient deprivation, because OSNs are the only mammalian sensory receptors that intrinsically regenerate throughout an animal’s life (Schwob, 2002; Cheetham et al., 2016), and regeneration rapidly restores peripheral input unless the stem cell population in the OE is also eliminated.

More duration-controlled approaches to olfactory deprivation require blocking airflow to the nostril rather than lesioning OSNs, either through surgical occlusion or cauterisation of one nostril (Baker et al., 1993), or through placement of nose plugs (Cummings and Brunjes, 1997). These plugs can be easily inserted into the nostrils of anaesthetised mice (or rats) to fully block nasal airflow and later removed to restore normal odour sampling (Coppola, 2012; Galliano et al., 2021; Byrne et al., 2022). While bilateral nostril closure is not feasible, as mice are obligate nose breathers and lack the capacity for oral respiration (Xie et al., 2015), unilateral naris occlusion enables controlled, side-specific sensory deprivation and is widely used to study activity-dependent changes in the olfactory bulb. In pre-weaning mice, nostril cauterisation followed by surgical reopening remains the standard unilateral naris occlusion approach (Fujimoto et al., 2023; Inoue et al., 2021), as nasal growth renders fixed-size plugs ineffective. In post-weaning mice and rats, by contrast, nose plugs provide a practical and easily implemented unilateral naris occlusion method. Here, we provide a detailed protocol for the manufacture and use of custom-made nose plugs, pioneered by Cummings and Brunjes (1997), including insertion, removal, verification of occlusion, and histological validation, based on this established approach and as implemented and validated in our laboratories.

## RESULTS

### Nose plug manufacturing

The nose plug is made of three elements: an approx. 5 mm-long piece of tygon tubing of the same diameter as the mouse nasal cavity, unscented dental floss, and non-absorbable suturing thread. The occlusion is achieved by enlarging and blocking the tube’s lumen by pulling inside the floss around which multiple overlapping surgical thread knots have been placed. This central swelling pushes against the tube walls, effectively blocking the airflow through its lumen (Fig. 1).

**Figure 1.**
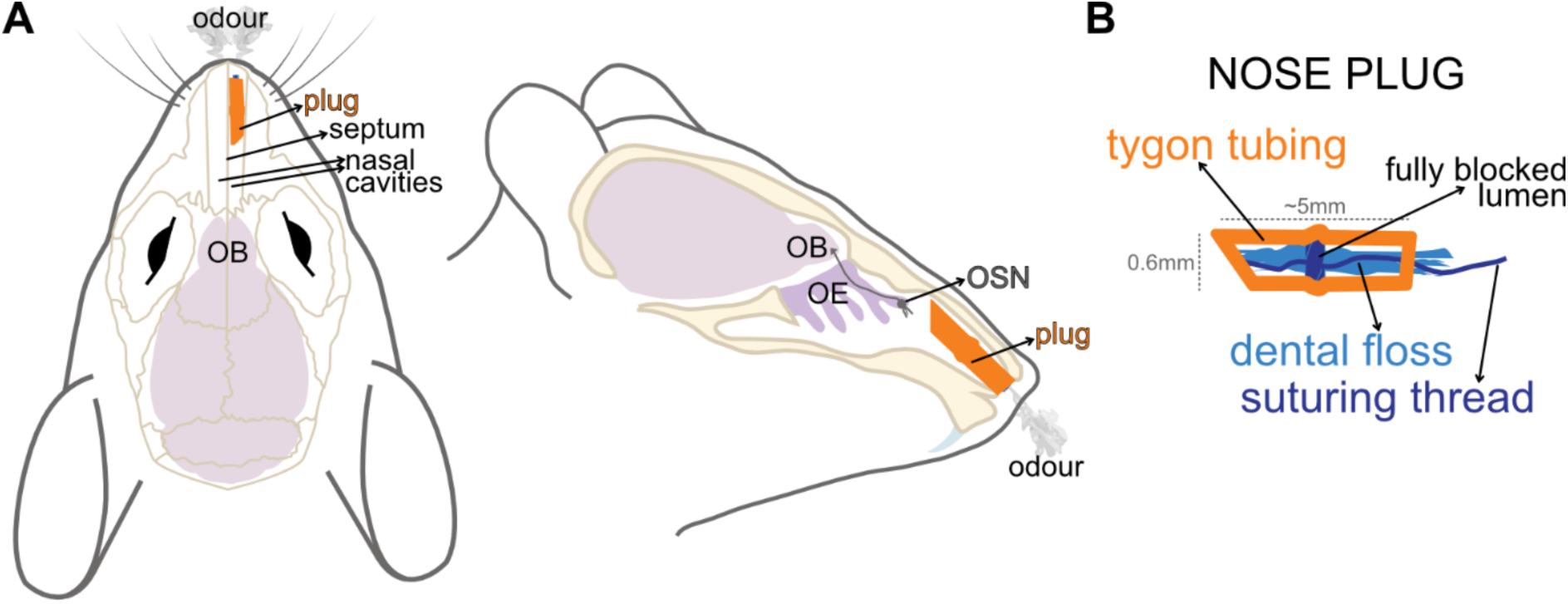
Schematic representation of the naris occlusion procedure. **(A)** Top and lateral view of the nose plug (orange), mouse skull (sepia), and brain (pink) inside the mouse head. Note that the two long nasal cavities are fully separated by a bone septum, effectively separating right and left sides. OB = olfactory bulb; OE = olfactory epithelium; OSN = olfactory sensory neuron. **(B)** Schematic representation of the custom-made nose plug, with the fully blocked tube lumen.

To start assembling the nose plug, the unscented dental floss needs to be broken down in thinner threads to achieve the appropriate thickness of floss that will fit inside the tubing and the nostril (note that while the tygon tubing is expandable and a full floss thread will fit, the nasal cavity is not). Using fine-pointed tweezers, flatten and separate the floss into separate strings to select either half (small plugs for juvenile mice up to P40) or 2/3 (bigger plugs for adult mice) of its width (Fig. 2A). Next, the suturing thread is knotted around the strand of the now separated dental floss strands, right in the middle (Fig. 2B, magenta arrow). Two further knots on top of the first knot are then secured, whilst leaving a long thread tail on one end. It is essential that the knots are made on top of each other (Fig. 2C’, magenta arrow), as they provide the needed volume to obstruct the tube and block the airflow. With sharp scissors, a piece of tygon tubing of an approximate 5 mm length is cut and the long tail of the suturing thread is pulled inside the tube’s lumen (Fig. 2C, green arrow). The long tail of the suturing thread is grasped with straight tweezers, with its end held between the tweezer’s arms. The thread tail is wrapped three times around the closed straight tweezers for a secure hold. Simultaneously, using the other hand, the middle of the tubing is grasped with blunt tweezers. The tubing is held firmly while the thread is pulled with steady, strong force towards the tubing. This will drag the knots and dental floss into the tube’s lumen, causing a bulge in the middle that effectively blocks its diameter.

**Figure 2.**
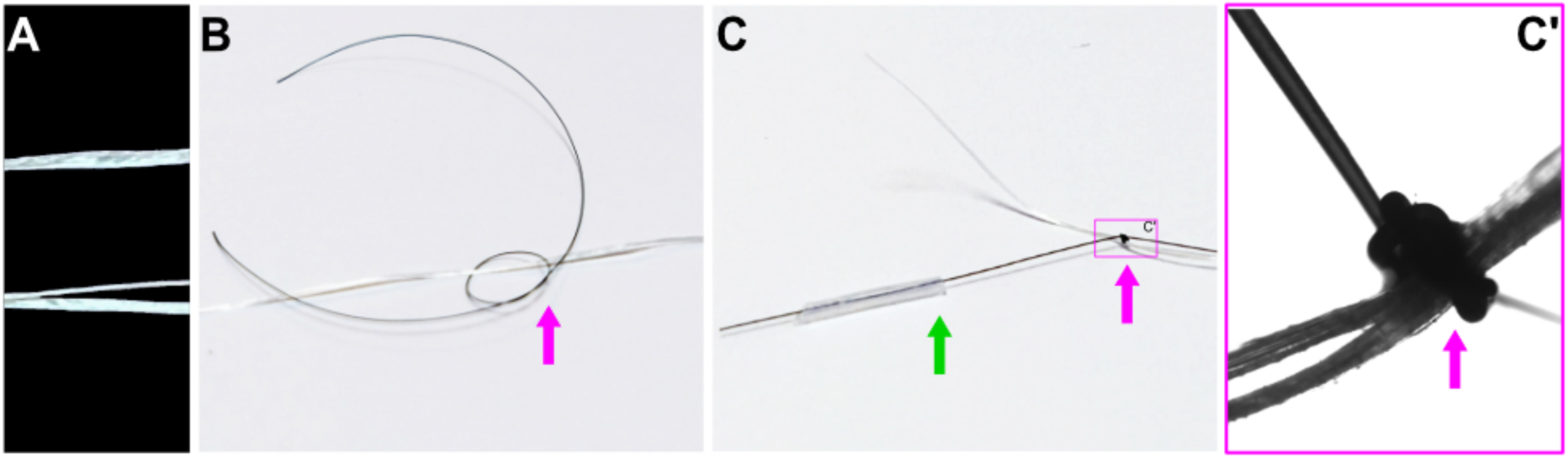
Selecting and knotting the floss and thread. **(A)** Example image showing a full floss string (top) and a floss string split into two halves (bottom). **(B)** Position of the first knot around the floss thread. The magenta arrow points at the future knot. **(C)** The three tight knots made around the floss (magenta arrow). One of the ends of the suture thread is placed inside the tube for the pulling step (green arrow). **(C’)** Bright-field microscope image of the knots and floss highlighted in C. The magenta arrow is pointing at the knot formation on top of each other which will result in occlusion.

To ease the insertion of the plug in the mouse nostril, it is essential to trim one end of the tube into a pointed, arrow-like shape (Fig. 3A, B). The cut must be done with either sharp scissors or a razorblade on the side without dental floss, as it has likely been damaged with the blunt tweezers in the previous step. A second cut will be performed on the opposite side of the tube to remove the excess of floss and suture thread whilst maintaining a short tail to facilitate visibility during storage and handling during insertion in the nostril (Fig. 3D-E). All plugs must be examined under a stereoscope, dissection microscope, or magnifying glass to confirm their suitability for occlusion. A good plug will be straight, have tightly secured knots that produce visible bulging of the tube, showing effective blocking (Fig. 3C, left). Plugs with loose knots should not be used as they may fail to achieve full occlusion (Fig. 3C, right).

**Figure 3.**
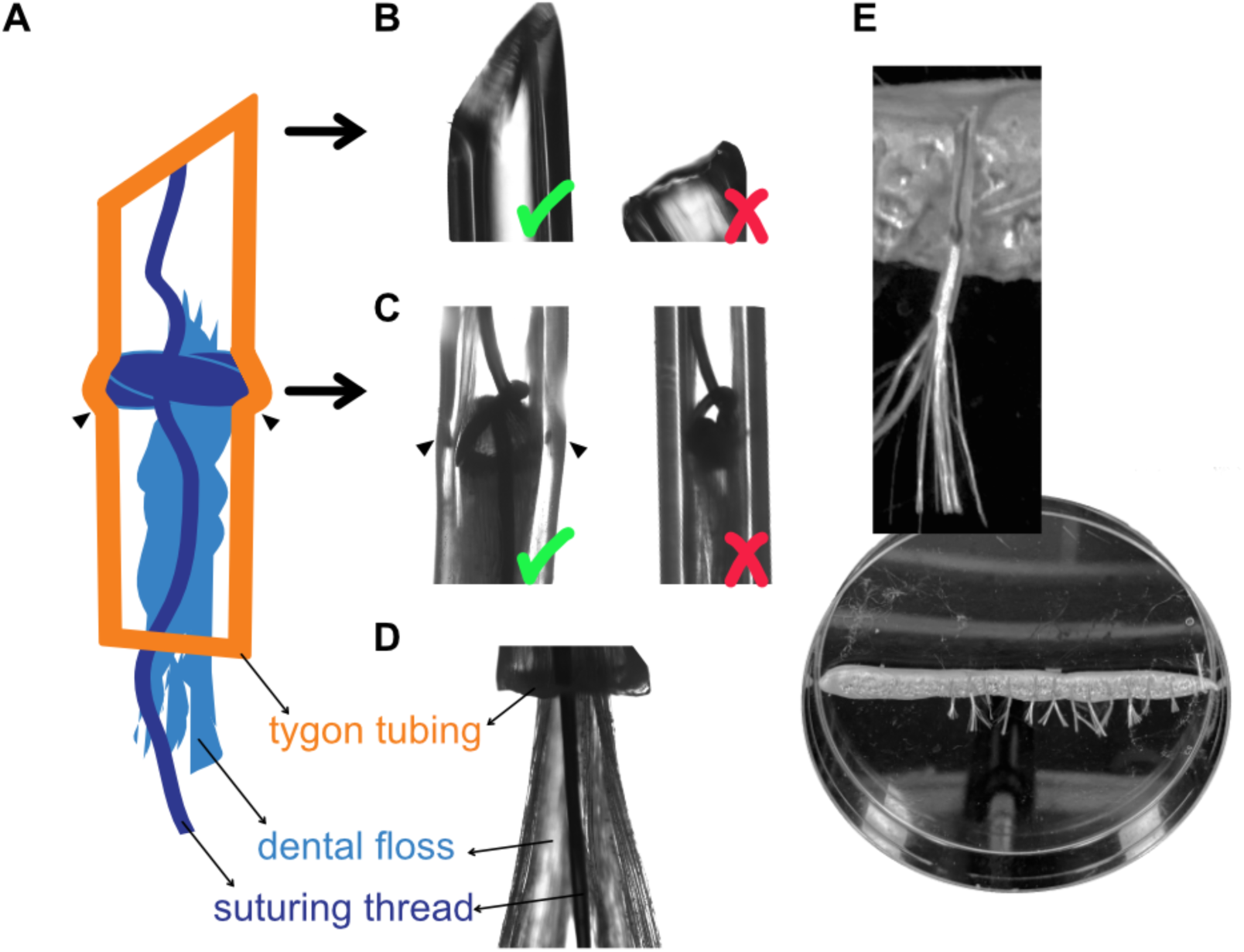
Checking and storing nose plugs. **(A)** Schematic drawing of the plug and **(B-D)** corresponding bright-field microscope images at the tip (cut vs uncut), middle (lumen blocked and tubing distended, triangles), and tail. Green ticks indicate the correct shape, red crosses wrong shapes. **(E)** Example images of plugs adhered to blue tack in a petri dish for storage. The excess of floss and suture thread facilitates easy visualisation and handling.

### Insertion of the nose plug

Mice must be briefly anaesthetised to insert and/or remove the nose plug to ensure a safe and stress-free procedure, and to allow for careful confirmation of plug placement.

#### Step 1: preparation of the workstation

The workstation needs to allow access to either a surgery microscope or a tabletop free standing magnifying glass with built in light (*e.g.* those used for soldering or crafts) and to an inhalation anaesthesia machine (recommended, *e.g.* isoflurane with induction box and scavenger unit). It is possible to use an injectable anaesthetic (*e.g*., ketamine/xylazine), which however will provide a much longer than needed length of anaesthesia, but which has the advantages of not requiring specialised equipment and of allowing more time for plug insertion. If inhalation anaesthesia with isoflurane is used, the entire procedure will take 5-10 minutes and therefore heating mats are not necessary, but if using injectable anaesthetics appropriate thermoregulation must be provided. Further preparation of the workstation includes draping the bench with tissue paper or monochromatic cloth on top of which the anaesthetised mouse will be placed (and on which will be easy to locate any accidentally-dropped plugs), and placing within reach of the lubricated plugs, the open petri dish with the spare pre-prepared plugs, blunt tweezers, and sharp scissors. The selected plugs are lubricated by placing the plug’s tip into the open unscented Vaseline jar, enough to facilitate insertion, but not make gripping the plug with tweezers difficult.

#### Step 2: isoflurane anaesthesia and plug insertion

The mouse is placed in the anaesthesia induction box, and after induction is achieved the isoflurane level is reduced to maintenance level. It is important to achieve deep, stable breathing for at least one full minute before removing the mouse from the induction and move it to the workstation. Leaving the isoflurane machine on with maintenance settings, the mouse is then placed on the workstation. At any point from this stage onwards, if the mouse starts giving clear signs of waking up, it must be promptly placed back in the anaesthesia box again and left there until deep stable breathing is once more achieved.

To begin the plug insertion, the fur on the mouse’s neck is gently but tightly grabbed to hold it sideways and the mouse positioned under the magnifying glass or microscope to visualise its nostrils. The grip should be strong enough to retract the snout and expose the nostril, while maintaining a stable side view to facilitate plug insertion. For right-handed experimenters, holding the mouse with the left hand, exposing its right nostril, and grabbing the plug with the right hand is recommended – and vice versa for the left-handed.

Using the blunt tweezers, the plug left ready in the Vaseline tub is securely held onto its ungreased half and inserted in the magnified mouse nostril. Important: the mouse nasal cavity curves upwards after the nostril opening, so to avoid hitting the nasal cavity floor the plug needs to be aimed with a slight upwards bend and not horizontally (Fig. 4A, B). If resistance is encountered, the plug must be removed, tilted upward further, and re-inserted: it should go in without force.

**Figure 4.**
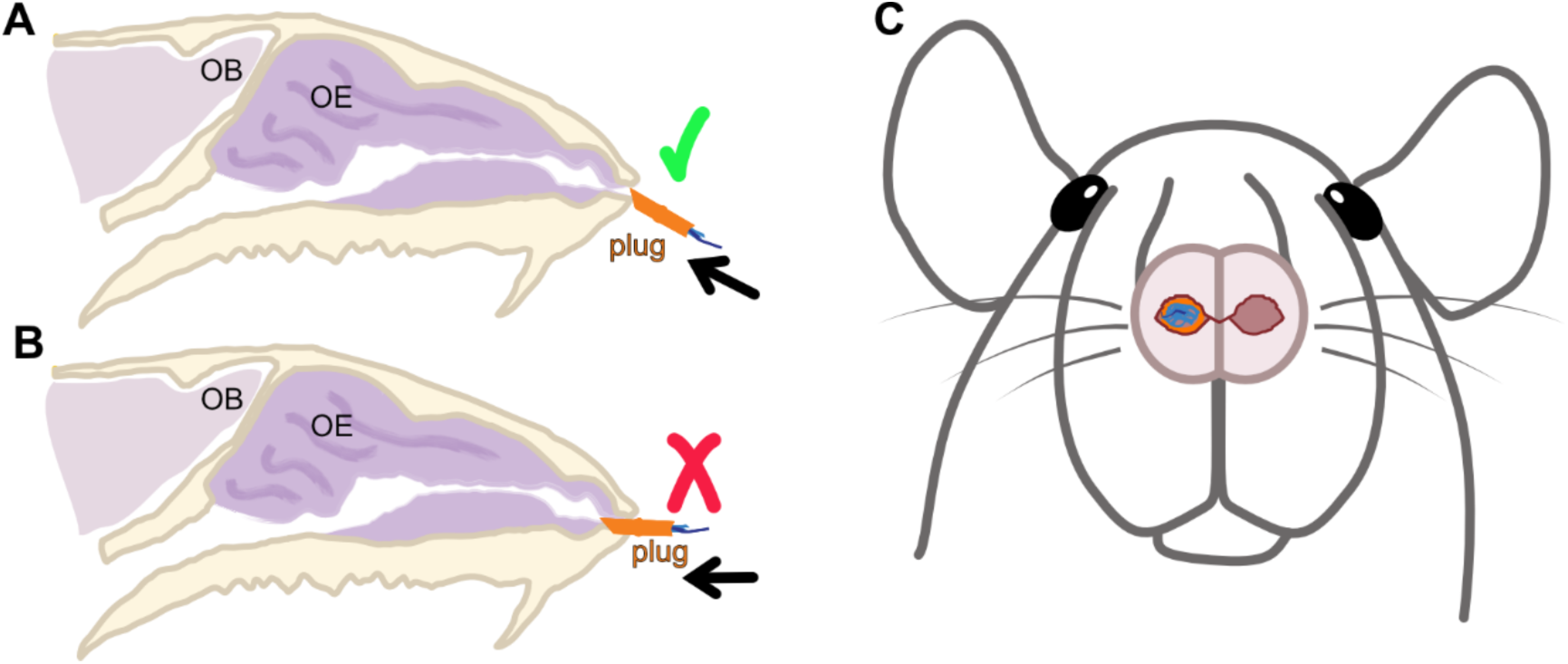
Inserting and trimming the plug. Schematic drawing of the correct **(A)** and wrong angle **(B)** at which the plug needs to be inserted in the mouse’s nostril. Note in B how by angling it straight, it hits the bone and misses the nasal cavity. **(C)** Schematic drawing of the mouse nostrils and right-inserted plug with enough suturing thread tail (dark blue) visible within the nasal cavity to allow for optional plug removal.

#### Step 3: nose plug trimming and mouse recovery

Once the plug is correctly inserted in the nostril, the mouse can be further rotated to a belly-up position to have a full visual of both nostrils under the microscope or magnifying glass (Fig. 4C). This will allow for careful confirmation of the plug placement, including the black tails of suturing thread being clearly visible but not hanging outside the nostril; use the sharp scissors and trim as required. This step is essential because the mouse, or its cage mate, can easily use hanging threads to pull the plug out. Especially if the experiment requires plug removal, it is crucial to find the right balance: leaving enough of the suturing thread “tail” inside the nostril for easy retrieval with pointy tweezers for experimental plug removal while ensuring that the mouse cannot remove it on its own. Once the experimenter is satisfied with the plug placement and trimming, the mouse is placed back in its home cage where it will wake up very rapidly (within 1-2 minutes) and start grooming its face around the nose. This is a good sign since it indicates that the plug is inserted. After a few minutes of attempted extraction, mice adapt and stop grooming the nose. If persistent grooming occurs and/or the mouse shows signs of distress or discomfort (never observed in our hands, but theoretically possible), humane endpoints outlined in the relevant ethical approval should be implemented. While monitoring the full recovery of the mouse, the workstation can be cleaned and the isoflurane scavenged from the anaesthesia box, and carefully written records of which nostril (right or left) has been plugged can be completed.

### Removal of the nose plug and optional plugging of the other nostril

One of the advantages of nose plugging versus nostril cauterisation is its ease of reversibility after a defined length of sensory deprivation. To remove the plug, the same workstation described above can be prepared, with the addition of a set of pointy straight tweezers. The deeply anaesthetised mouse is once again held by the skin on its neck and turned belly-up under the microscope or magnifying glass to locate the plug’s black suturing thread tail (Fig. 4C). Using point tweezers while making sure that tissue is not pricked, the suturing thread tail is grabbed firmly, and the plug is pulled out of the nose. The mouse is then allowed to wake up in its home cage or alternatively, a new plug can first be inserted in the previously unplugged nostril.

### Confirmation of plug location at the end of the experiment

At the end of the naris occlusion experiment, it is essential to confirm that the plug remains in place within the nasal cavity. This is necessary because the plug may fall out if not inserted correctly, and although rare, animals may also remove it despite correct placement (in our experience, this occurred in <5% of procedures performed by trained experimenters). If the plug is not present in the nostril, the time of removal cannot be determined, and the data must therefore be excluded. When plug removal is included as part of the protocol, this step inherently provides confirmation of plug retention. In cases where the animal is culled without prior plug removal, the nasal cavity should be dissected to visually confirm that the plug remains in the correct nasal passage. Care must be taken during dissection of the nasal bones to avoid damaging or cutting the plug, which would preclude confirmation of its placement.

### Assessing the success of olfactory deprivation

The success of olfactory deprivation can be evaluated through a wide range of experimental approaches, as naris occlusion at different ages and durations induces substantial changes in OSNs and OB neurons identified using electrophysiological recordings and imaging in slices and *in vivo* (Tyler et al., 2007; He et al., 2012; Kass et al., 2013), as well as modifications in gene expression profiles in OSNs, detected by quantitative PCR and *in situ* hybridisation (Coppola and Waggener, 2012; Fischl et al., 2014; Tufo et al., 2025). Numerous previous studies have consistently shown that naris occlusion reduces the expression of the neuronal activity marker cFos and the rate-limiting enzyme in the dopamine synthesis pathway tyrosine hydroxylase (TH) in the dopaminergic neurons of the ipsilateral bulb to the closed nostril, with significant changes detectable as early as 1 day after occlusion (Baker et al., 1993; Liu et al., 1999; Galliano et al., 2021; Byrne et al., 2022). The expression of these activity markers can be readily assessed using immunohistochemistry on brain slices, a straightforward, widely accessible, and highly reproducible approach. However, accurate comparison of staining intensity requires careful control of variability arising from antibody binding, tissue penetration, and processing conditions (Vlug et al., 2005). To address this, OBs from occluded and control animals should be co-embedded in agarose blocks to generate matched “sets”, which are then sliced using a vibratome. Multiple slices from each set ideally spanning the rostro-caudal axis of the OB, can then be processed and imaged together under identical conditions. For each slice within a set, TH and cFos staining intensities should be normalised to the mean intensity of control OBs present in that same slice, enabling reliable comparisons while minimising technical variability (Galliano et al., 2021).

Using the naris occlusion protocol described above, which was performed independently by two experimenters in two laboratories (Cambridge and London) over a three year period, with occlusion maintained for one day, we consistently observed marked reductions in both TH (Fig. 5B; Mean ± SEM, Linear mixed-effects model nested on mouse, effect on treatment; Cambridge TH: Sham Control = 100.0 ± 1.4 %, n=783, N=3; Occluded = 92.8 ± 1.3 %, n=843, N=3. (β = −7.20, SE = 1.88, t(1624) = −3.83, p = 1.3 × 10^-4^; Type III ANOVA: F(1, 1624) = 14.65, p = 1.3 × 10⁻⁴); London TH: Sham Control = 100.0 ± 1.0 %, n=1513, N=3; Occluded = 76.6 ± 0.9 %, n=1745, N=3. (β = −0.23, SE = 0.064, t(4.03) = −3.54, p = 2.37 × 10^-2^; Type III ANOVA: F(1, 4.03) = 12.546, p = 2.37 × 10^-2^)) and cFos (Fig. 5D; Mean ± SEM, Linear mixed-effects model nested on mouse, effect on treatment; Cambridge cFos: Sham Control = 97.6 ± 2.7 %, n=795, N=3; Occluded = 67.7 ± 1.5 %, n=879, N=3. (β = −32.346, SE = 8.592, t(3.85) = −3.765, p = 2.1 × 10^-2^; Type III ANOVA: F(1, 3.85) = 14.175, p = 2.1 × 10^-2^); London cFos: Sham Control = 100.0 ± 1.0 %, n=1513, N=3; Occluded = 56.4 ± 0.6 %, n=1745, N=3. (β = −0.44, SE = 0.053, t(4.03) = −8.26, p = 1.13 × 10^-3^; Type III ANOVA: F(1, 4.03) = 68.162, p = 1.13 × 10^-3^)) expression across experiments. This was further strengthened by the use of two different cFos antibodies (see methods) under otherwise identical experimental conditions, demonstrating the reliability of this approach (Fig. 5C, E; Two-way ANOVA manipulation with Location, Treatment and their interaction as factors; TH intensity Treatment *F*(1,8) = 21.588, *p* = 1.65 × 10^-3^, Location *F*(1,8) = 4.728, *p* = 6.1× 10^-2^, Location*Treatment *F*(1,8) = 4.728, *p* = 6.1× 10^-2^; cFos intensity Treatment F(1,8) = 56.42, *p* = 6.85× 10^-5^, Location (F(1,8) = 0.327, *p* = 0.58, Location*Treatment *F*(1,8) = 1.245, *p* = 0.297). Our findings are also consistent with our previous studies reporting similar TH and cFos downregulation following one day of sensory deprivation (Galliano et al., 2021; Byrne et al., 2022).

**Figure 5.**
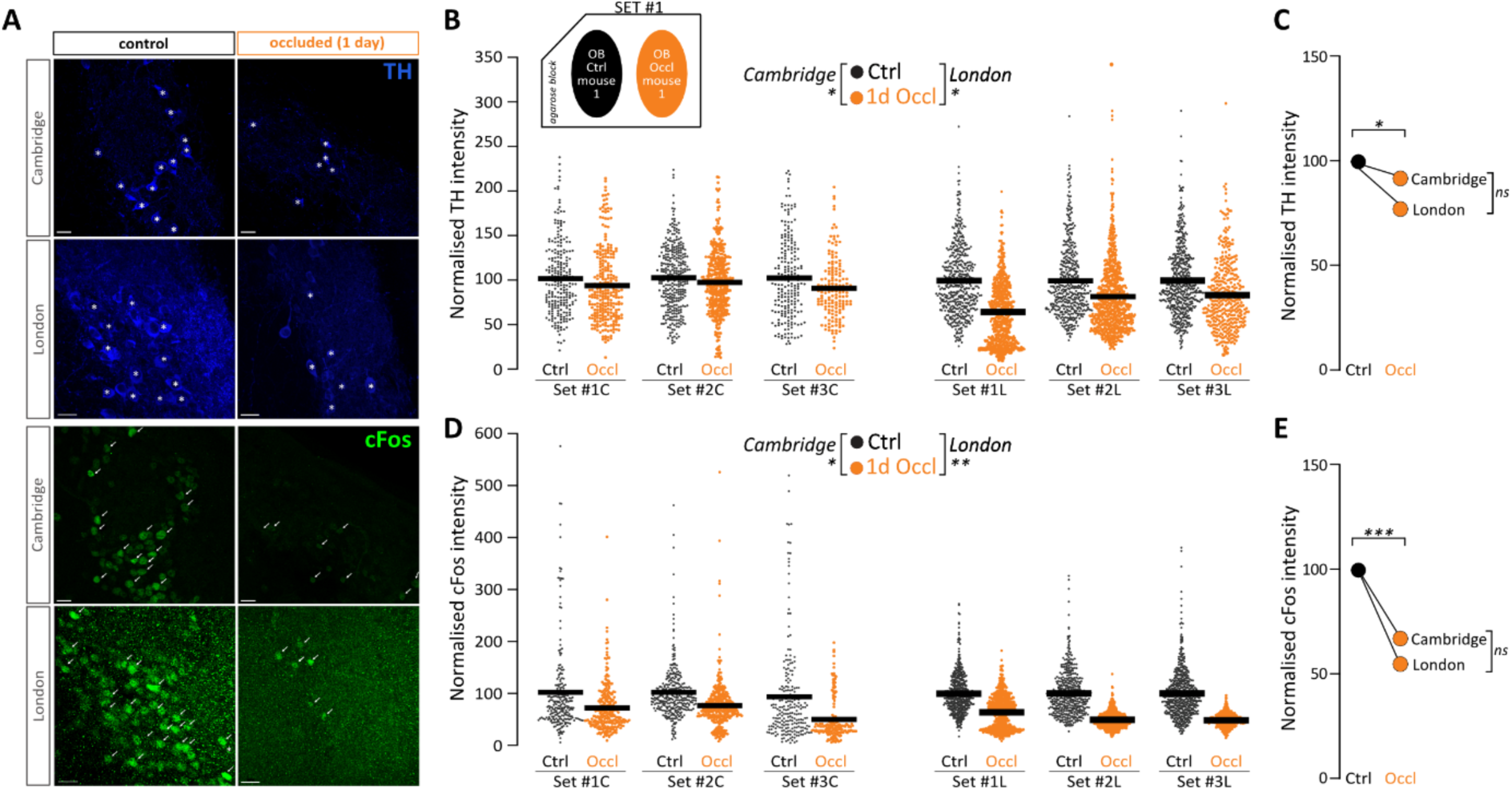
Assessing naris occlusion efficiency via TH and cFos staining across labs and experimenters. **(A)** Example images of tissue from sham control and 1-day occluded mice from independent experiments conducted at the University of Cambridge (C) and at King’s College London (L) showing bulbar dopaminergic neurons stained with TH (top) and cFos (bottom). The asterisks show examples of TH+, the arrows of cFos+ cells in both groups. **(B)** TH intensity levels normalised to co-embedded controls in three “sets” (see insets) in sham control and 1-day occluded mice. Small circles are individual datapoints, horizontal black lines are mean, each column is a mouse, each set is a pair of co-embedded control mouse and occluded mouse. **(C)** Mean normalised TH intensity across the three sets quantified across the two locations. **(D-E)** Same conventions as in B-C for cFos quantification in bulbar dopaminergic neurons. Scalebars are 20 μm; * p<0.05; ** p<0.01; *** p<0.001.

A major caveat of this type of analysis is that the same activity-dependent marker (TH) is used both to identify dopaminergic neurons and to quantify the success of the manipulation. Moreover, dopaminergic neurons exhibit adult neurogenesis (Bonzano et al., 2016), and prolonged sensory deprivation could potentially influence the maintenance and turnover of this population in an activity-dependent manner (Angelova et al., 2023). To address this, we validated the use of TH immunoreactivity as a measure of sensory deprivation-induced changes using a longer occlusion paradigm (one week) in a transgenic mouse line in which dopaminergic neurons are genetically labelled by tdTomato expression under the DAT-Cre driver line (Bäckman et al., 2006) and are also immunostained for TH (Fig. 6A). We first confirmed that DA cell density was unchanged following 7 days of occlusion (Fig. 6B; Mean ± SEM; Sham Control = 124.12 ± 10.49, n=21, N=5; Occluded = 131.99 ± 13.22, n=21, N=5. Linear mixed-effects model nested on mouse, effect on treatment (β = 11.45, SE = 34.70, t(7.9) = 0.33, p = 0.750; Type III ANOVA: F(1, 7.9) = 0.1089, p = 0.750). We then quantified TH expression in sham control and 7-day occluded OBs in dopaminergic neurons identified either by TH immunoreactivity or by genetically encoded tdTomato labelling. In both cases, neurons from occluded mice exhibited reduced TH expression, demonstrating that both approaches provide valid strategies for identifying dopaminergic neurons and assessing sensory deprivation-induced changes in TH levels. The treatment effects were significant for both identification strategies (Fig. 6C; TH-identified neurons: β = −33.24, SE = 4.39, t(8.16) = −7.58, p = 5.82 × 10⁻⁵, sham n=2161, occluded n=1563; tdTomato-identified neurons: β = −26.20, SE = 7.00, t(8.07) = −3.75, p = 0.00557, sham n=1609, occluded n=1437; N=5 mice per group for both analyses). However, reduced DA neuron density has been reported following 7 days of occlusion (Grier et al., 2016), and even longer periods of sensory deprivation may induce additional changes in DA neuron maintenance or population dynamics (Baker et al., 1993; Lazarini et al., 2014; Angelova et al., 2023; Huang et al., 2024). Therefore, the use of a non-activity-dependent marker to identify these neurons is strongly encouraged.

**Figure 6.**
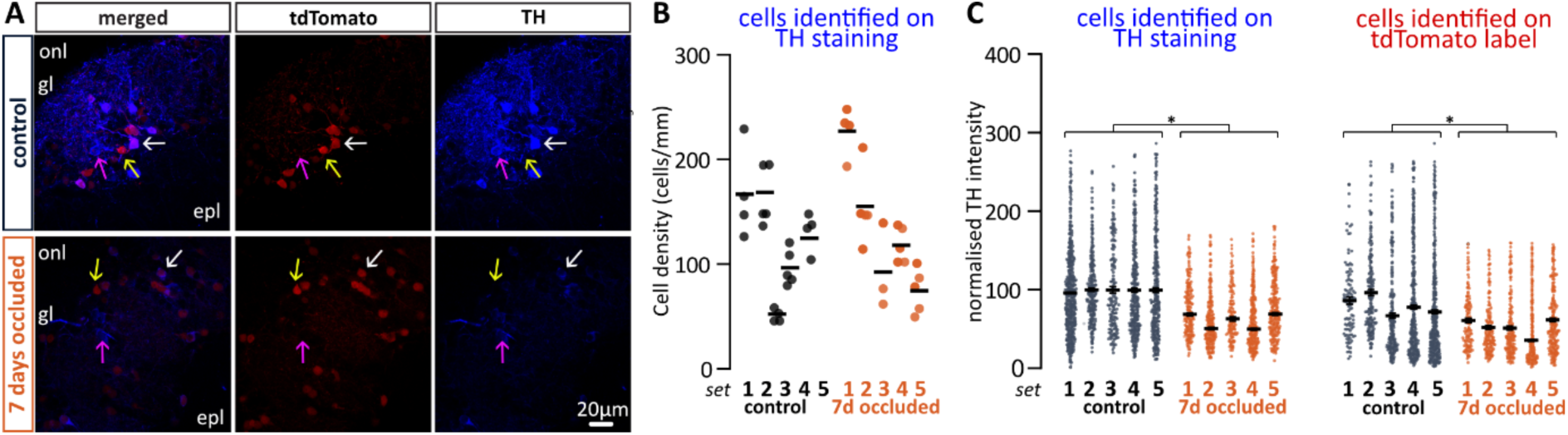
Validation of TH immunoreactivity as a measure of longer-lasting sensory deprivation-induced changes in DA neurons. **(A)** Representative images of olfactory bulbs from DAT-Cre::tdTomato mice subjected to sham control or 7-day naris occlusion, showing genetically labelled dopaminergic neurons (tdTomato) and TH immunoreactivity**. (B)** Quantification of dopaminergic neuron density following 7 days of naris occlusion. **(C)** Quantification of TH expression levels in DA neurons identified by TH immunoreactivity or tdTomato labelling. In both cases, DA neurons from occluded animals exhibited reduced TH expression compared with controls. Small circles are individual stacks (B) or datapoints (C), horizontal black lines are mean, each column is a mouse, each set is a pair of co-embedded control mouse and occluded mouse; scalebars are 20 μm; * p<0.05.

### Evaluation of the contralateral bulb (free nostril side) as an intra-animal control

To evaluate the suitability of the contralateral free nostril side as an intra-animal control (Tepe et al., 2018), we applied our co-embedding strategy and analysed both olfactory bulbs rather than restricting analysis to a single bulb per mouse (Fig. 7A). This allowed direct comparison of unperturbed and sham-occluded controls, as well as occluded and contralateral free nostril bulbs from the same animal, following either 1 day or 7 days of naris occlusion.

**Figure 7.**
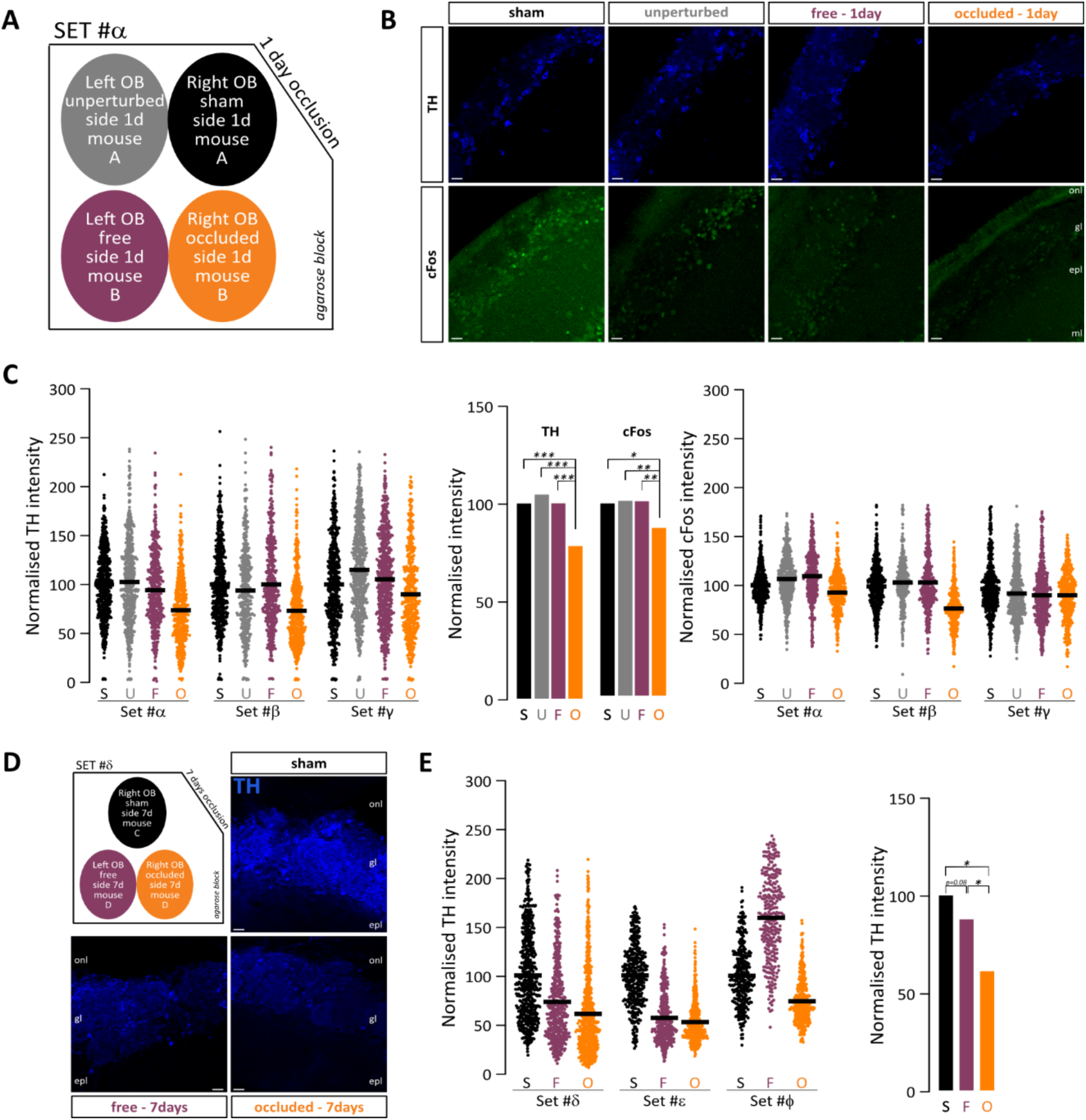
Evaluation of the contralateral free-side olfactory bulb as an intra-animal control following sensory deprivation. **(A)** Schematic representation of the co-embedding strategy, in which each agarose block contained four OBs: the left and right OBs from a control mouse which received a sham plug in the right nostril, and the left and right OBs from a mouse whose right nostril had been occluded for 24 h. **(B**) Representative images of OB tissue from sham control and 1-day occluded mice stained for TH (top) and cFos (bottom). **(C)** TH and cFos intensity normalised to the co-embedded sham control, across three sets of samples, comprising sham control (black), unperturbed (grey), free nostril (purple), and occluded nostril (orange) conditions. Small circles represent individual data points; horizontal black lines indicate the mean. Each column represents one mouse, and each set comprises a control mouse and an occluded mouse whose OBs were co-embedded. Bar plots show mean normalised TH and cFos intensity across the three sets**. (D–E)** Representative images and quantification of TH immunoreactivity in bulbar dopaminergic neurons from a sham control mouse and from the free and occluded sides of a mouse whose right nostril had been occluded for 7 days, following the same conventions as in (B–C). Scalebars, 20 µm; onl = olfactory nerve layer, gl = glomerular layer, epl = external plexiform layer; *p<0.05; **p<0.01; ***p<0.001.

First, we confirmed that unperturbed and sham-occluded animals exhibited equivalent TH and cFos expression (Fig. 7; Mean ± SEM, Generalised linear mixed-effects model nested on mouse, effect on condition, contrasts corrected using two stage Benjamini-Krieger-Yekutieli; TH: Sham Control = 100.0 ± 0.95%, n=1744, N=3; Unperturbed = 104.7 ± 1.09%, n=1697, N=3; Sham-Unperturbed(β = −0.03, SE = 0.05, z = −0.562, p = 0.526); cFos: Sham Control = 98.5 ± 0.56 %, n=1721, N=3; Unperturbed = 99.91 ± 0.66%, n=1613, N=3; Sham-Unperturbed(β = −0.02, SE = 0.048, z = −0.509, p = 0.736)), indicating that either approach provides a suitable control condition (Fig. 7; Mean ± SEM, Generalised linear mixed-effects model nested on mouse, effect on condition, contrasts corrected using two stage Benjamini-Krieger-Yekutieli; TH: Occluded = 78.31 ± 0.96 %, n=1619, N=3; Sham-Occluded(β = 0.21, SE = 0.05, z = 4.1, p = 6.55×10^-5^), Unperturbed-Occluded(β = 0.24, SE = 0.05, z = 4.6, p = 2.14×10^-5^); cFos: Occluded = 86.11 ± 0.57 %, n=1619, N=3; Sham-Occluded(β = 0.13, SE = 0.05, z = 2.49, p = 0.017), Unperturbed-Occluded(β = 0.16, SE = 0.05, z = 2.9, p = 0.008)). Following 1 day of occlusion, the contralateral free nostril bulb was also indistinguishable from independent control conditions (Fig. 7; Mean ± SEM, Generalised linear mixed-effects model nested on mouse, effect on condition, contrasts corrected using two stage Benjamini-Krieger-Yekutieli; TH: Free = 100.2 ± 1.04 %, n=1708, N=3; Sham-Free(β = 0.002, SE = 0.05, z = 0.03, p = 0.951), Unperturbed-Free(β = 0.03, SE = 0.05, z = 0.573, p = 0.526), Free-Occluded(β = 0.210, SE = 0.05, z = 4.163, p = 6.55×10^-5^); cFos: Free = 99.63 ± 0.70 %, n=1666, N=3; Sham-Free(β = −0.014, SE = 0.05, z = −0.258, p = 0.736), Unperturbed-Free(β = +0.01, SE = 0.05, z = 0.192, p = 0.736), Free-Occluded(β = 0.148, SE = 0.04, z = 3.292, p = 6 ×10^-3^)).

After 7 days of occlusion, expression of TH was similarly reduced in occluded bulbs compared to those from sham-occluded animals (Fig. 7; Mean ± SEM, Generalised linear mixed-effects model nested on mouse, effect on condition, contrasts corrected using two stage Benjamini-Krieger-Yekutieli; Sham Control = 100.0 ± 1.03%, n=1349, N=3; Occluded = 61.57 ± 0.77 %, n=1812, N=3; Sham-Occluded(β = 0.357, SE = 0.15, z = 2.307, p = 0.0337). However, the bulbs from the free nostril exhibited an intermediate phenotype very inconsistent across the three sets (Fig. 7; Mean ± SEM, Generalised linear mixed-effects model nested on mouse, effect on condition, contrasts corrected using two stage Benjamini-Krieger-Yekutieli; Free = 87.93 ± 1.48%, n=1346, N=3; Sham-Free(β = 0.193, SE = 0.16, z = 1.203, p = 0.0826); Free-Occluded(β = 0.164, SE = 0.07, z = 2.295, p = 0.0337).

Thus, while expression on the free side was consistent across biological replicates after 1 day of occlusion, the marked inter-replicate variability observed after 7 days, ranging from levels comparable to the occluded side to above control levels, highlights the potential limitations of using the free side as an intra-animal control following prolonged sensory deprivation.

## DISCUSSION

In this methods article, we provide a standardised and validated protocol for inducing reversible unilateral sensory deprivation in juvenile and adult mice using custom-made naris plugs. Building on previously established approaches (Cummings and Brunjes, 1997; Coppola, 2012), we describe in detail the construction of reliable plugs using readily available commercial materials and provide practical guidance to improve reproducibility across laboratories. We further outline strategies for consistent quantitative validation of deprivation-induced changes, including controlled tissue processing and immunohistochemical analysis using matched samples. The successful implementation of this protocol across independent laboratories using only the instructions provided here demonstrates its robustness and accessibility, establishing a practical framework for the reproducible deprivation of olfactory sensory input.

### Advantages of performing naris occlusion using custom-made nose plugs

Naris occlusion is a powerful approach to manipulate sensory activity in the olfactory system and study experience-dependent plasticity. By reducing sensory input in a controlled and reversible manner, it enables direct investigation of how olfactory circuits respond to decreased activity, providing a more defined experimental manipulation than approaches that increase sensory experience, such as environmental enrichment. Importantly, the effects of sensory deprivation can be directly validated using molecular and cellular readouts, such as TH and cFos, allowing experimental manipulation and its consequences to be quantitatively assessed (Baker et al., 1993; Galliano et al., 2021; Byrne et al., 2022).

One of the main strengths of this procedure when performed with custom made nose plugs is its high reliability. While some nose plugs may be lost during the process, this occurs in only a small percentage of cases (<5%) and there is no ambiguity whether the plug was kept or lost since its presence can be readily confirmed at the end of the experiment. Most importantly, where the plug is retained, the protocol consistently induces decreased TH and cFos immunofluorescence, which have traditionally been used as activity readouts (this paper, (Kosaka et al., 1987; Baker et al., 1993; Wilson and Sullivan, 1995; Philpot et al., 1997; Briñón et al., 2001; Yan et al., 2007; Parrish-Aungst et al., 2011; Sawada et al., 2011; Lau and Murthy, 2012; Mast and Fadool, 2012; Bovetti et al., 2013; Chand et al., 2015; Galliano et al., 2021; Byrne et al., 2022)).

This high reliability is further enhanced by the other key advantages of the protocol. It is reversible, enabling temporary manipulation of the olfactory network to study cellular and circuit recovery after sensory deprivation, something that is not achievable with permanent manipulations. Additionally, the procedure is minimally invasive and ensures animal welfare: mice are only briefly anaesthetised, and insertion of the plug is quick and straightforward. Because the plug remains at the periphery of the nostril and does not contact internal structures, the procedure is well tolerated. There is also no need for specialised equipment, as the materials required for plug production and the experiment are commercially available and low-cost.

### Unperturbed, sham occluded, or free nostril: what to use as a control?

Mice use their nose not only for odour sampling but also for respiration, with important consequences when one nostril is occluded. Blocking one nostril disrupts normal bilateral airflow patterns and increases airflow through the open nostril, potentially creating a condition that combines altered odour exposure, modified airflow dynamics, and possible mechanical stress on the sensory epithelium (Maruniak et al., 1989; Coppola and Waggener, 2012; Kass et al., 2013; Chiesa Estomba et al., 2015; Weitzman et al., 2020). This raises an important methodological question: can the contralateral, non-occluded OB be considered a true intra-animal control? On one hand, using the ipsilateral to open nostril OB as a control provides the advantage of a within-animal comparison, reducing biological variability (Tepe et al., 2018; George et al., 2022). On the other hand, increased airflow through the open nostril could alter sensory input, potentially leading to either enhanced odour stimulation, mechanical effects, or a combination of both. Thus, the open nostril may not represent a completely unperturbed state. Alternative approaches include the use of unperturbed littermate controls (Galliano et al., 2021) or sham-occluded animals, in which a piece of the plug material is inserted and immediately removed, controlling for anaesthesia and handling while avoiding prolonged changes in airflow (Byrne et al., 2022). Although these approaches provide cleaner experimental controls, they require additional animals and do not allow within-animal comparisons. Our findings indicate that the contralateral open nostril may serve as a practical intra-animal control for short-term occlusion experiments based on the parameters examined here. However, the possibility that more sensitive measures, such as changes in neuronal firing properties, synaptic connectivity, or other aspects of circuit function (Huang et al., 2024), may reveal differences between the open nostril and a fully unperturbed control even at this short occlusion duration cannot be excluded. Moreover, performing statistical analysis using the free side as the control should account for repeated measures, as the source for both control and treatment is the same biological unit, the animal, negating any assumption of independence. Alternatively, a mixed-effects design that models animals as a source of correlation can be used. Restricting analysis to within-animal comparisons also complicates translating immunohistochemical or molecular findings to behaviour, because olfactory behaviour is difficult to assess in a side-specific (within-animal) manner. Therefore, independent unperturbed or sham-occluded littermate controls represent good experimental practice and are recommended, particularly for prolonged deprivation paradigms or experiments requiring strict control of sensory input.

### Limitations and challenges

While the plug-based occlusion method is highly effective for inducing sensory deprivation in mice, it has limitations. First, the procedure requires careful attention to plug preparation. When learning to make plugs, considerable practice and trial and error may be required to achieve the optimal fit. Second, although plug loss occurs only in a small proportion of animals, retention cannot always be controlled even when the plug has been properly inserted. Therefore, plug presence should always be verified at the end of the experiment, and animals in which the plug has been lost should be excluded from further analysis. Third, this protocol is applicable only to juvenile and adult mice and cannot be used in neonates since their noses are still growing. The plug, which remains the same size, becomes ineffective in occluding the nostrils as the nose develops. Consequently, researchers working with neonates often choose cauterisation as the method to induce naris occlusion, even though it is irreversible (Fujimoto et al., 2023). Fourth, plug-based occlusion does not allow for the occlusion of both nostrils given that mice are obligate nose breathers, but this is also a limitation of cauterisation. If complete olfactory deprivation is required, for instance to study behaviours such as navigation while ruling out olfactory contributions, this can be achieved through chemical lesions. However, olfactory sensory neurons regenerate (Cheetham et al., 2016), making this approach less suitable for long-term deprivation studies.

Finally, a further consideration when using immunohistochemistry as a quantitative readout and/or to validate the success of the occlusion, with plugs, with cauterization, or with chemical methods, is the need to control both biological and technical sources of variability. Differences between animals (Figs. 5–6–7), as well as variability introduced during tissue processing, antibody binding, staining, and imaging, can influence measured staining intensity. Matched co-embedding of control and occluded samples provides a simple strategy to minimise these effects by allowing tissues to be processed, stained, and imaged under identical conditions. Normalisation within matched tissue sets therefore improves the reliability of quantitative comparisons.

### Conclusions

In conclusion, this protocol provides a simple, reproducible, and accessible approach for inducing reversible unilateral sensory deprivation in adult and juvenile mice. By combining reliable naris occlusion with controlled validation strategies and appropriate consideration of experimental controls, this method enables robust investigation of activity-dependent plasticity in the olfactory system across different laboratories and experimental settings.

## MATERIALS AND METHODS

### Animals

We used mice of either sex and housed them under a 12h light-dark cycle in an environmentally controlled room with free access to water and food. WT C57BL/6 mice (Charles River) were used either as experimental animals, or to backcross each generation of transgenic animals DATIREScre (B6.SJL-Slc6a3tm1.1(cre)Bkmn/J, Jax stock #006660) and Ai9 (B6.Cg–Gt(ROSA) 26Sortm9(CAG-tdTomato)Hze/J; Jax stock #007909). All experiments were performed after postnatal days (P) 21 at King’s College and the University of Cambridge in accordance with the United Kingdom Animals (Scientific Procedures) Act 1986, under the auspices of Home Office personal and project licences held by the authors.

### Sensory deprivation

Nose plugs were constructed as described in the Results section using the materials listed in Fig. 8. High magnification images of nose plugs were acquired using a Thermofisher EVOS M5000 microscope, low magnification plug photos were taken using a Sony Xperia 10 IV camera.

**Figure 8.**
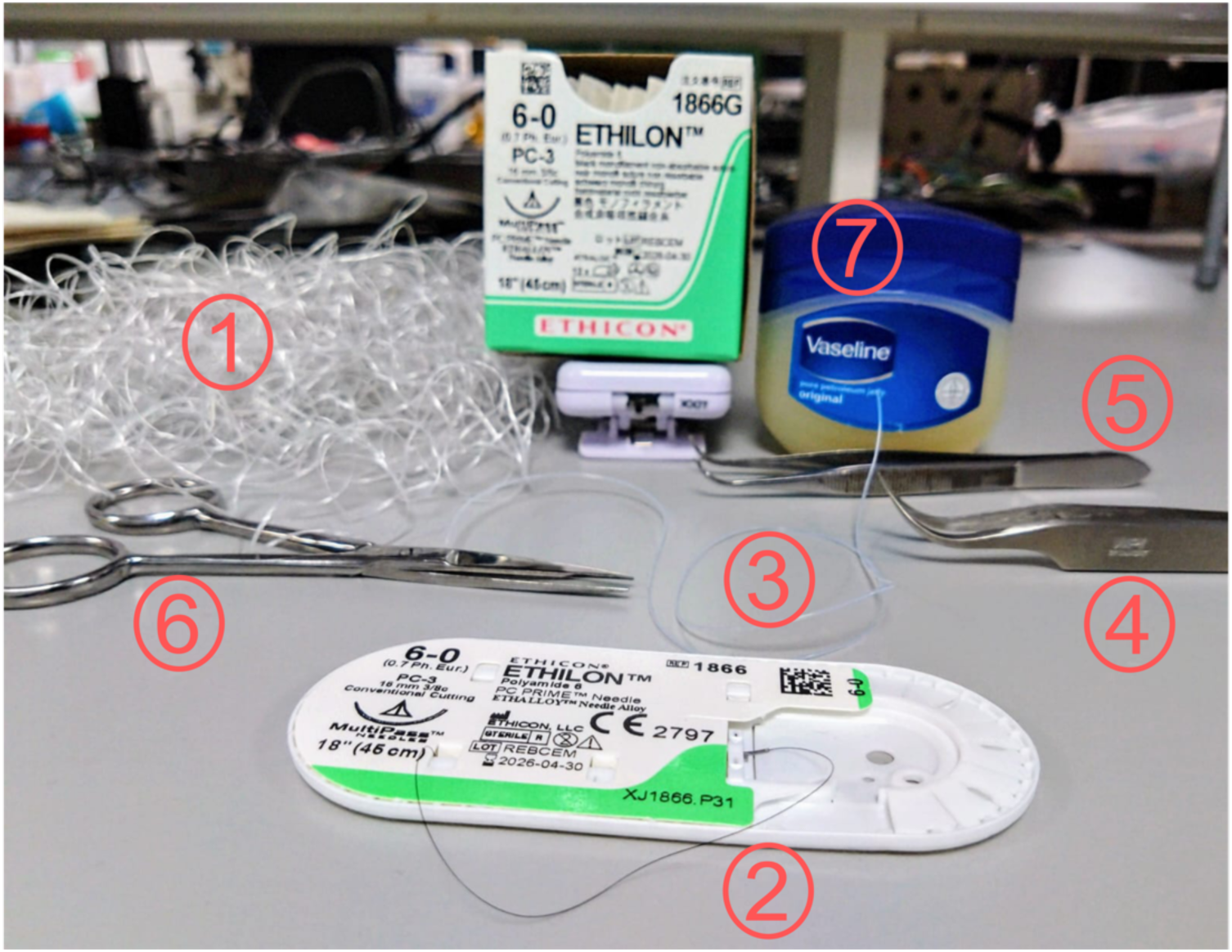
Nose plug preparation materials. (1) Unscented dental floss, removed from case and left to air dry for a few days. (2) Ethilon polyamide nonabsorbable suture, size 6-0 (Ethicon; UK supplier: AHP medicals). (3) tygon tubing with an outer diameter of 0.6 mm and inner diameter of 0.3 mm (VWR, #S1810-04). (4) Pointy curved and/or straight tweezers. (5) Blunt straight tweezers. (6) Sharp small scissors. (7) Unscented Vaseline.

Mice were briefly anaesthetised with isoflurane to allow insertion of a nose plug coated with Vaseline into either the right or left nostril. The plug remained in place for either 1 or 7 days. For experiments shown in Figures 5–6, only the olfactory bulb ipsilateral to the plugged nostril was used. For experiments shown in Figure 7, both ipsilateral and contralateral olfactory bulbs were analysed. Following transcardial perfusion, the nasal cavity was visually inspected to confirm that the plug remained intact. Animals in which the plug had been lost were excluded from subsequent analyses.

### Immunohistochemistry

Mice were anaesthetised with an overdose of pentobarbital and transcardially perfused with 15–20 ml of 1× phosphate-buffered saline (PBS) containing heparin (20 units/ml), followed by 15–20 ml of 1–4% paraformaldehyde (PFA; P001, TAAB Laboratories) prepared in a solution containing 3% sucrose, 60 mM PIPES, 25 mM HEPES, 5 mM EGTA, and 1 mM MgCl₂ (all reagents purchased from Sigma-Aldrich). The OBs were dissected and post-fixed in 1% PFA for up to 5 days. OBs ipsilateral to the plugged or sham-plugged nostril (or both bulbs from control and occluded mice) were co-embedded in a single 5–7% agarose block (“set”) and sectioned coronally at 50 µm using a vibratome (VT1000S, Leica, or 5100mz, Campden Instruments). For each experiment, a minimum of 3 sets (3 control, 3 occluded mice) was prepared. For each set, three or four sections were sampled across the rostrocaudal axis and processed for staining. Free-floating 50 µm sections were permeabilised and blocked with 5% normal goat serum (NGS; G9023, Sigma-Aldrich) in PBS/Triton/azide solution (0.25% Triton X-100 and 0.02% sodium azide) for 2 h at room temperature (RT). Sections were then incubated with primary antibody solution (antibodies diluted in 5% NGS in PBS/Triton/azide) for 2 days at 4°C. Primary antibodies used were mouse anti-tyrosine hydroxylase (TH; 1:500; Merck, MAB318) and rabbit anti-cFos (1:1000; Merck, ABE457, or 1:500; Cell Signaling, 2250). Following three 5-min washes in PBS, sections were incubated with species-appropriate Alexa Fluor-conjugated secondary antibodies (Life Technologies; diluted 1:1000 in PBS/Triton/azide) for 2 h at RT. After three additional 5-min washes in PBS, sections were mounted onto glass slides (J2800AMNZ, Epredia) using FluorSave (345789, Merck) or MOWIOL-488 (475904, Merck). For all experiments, staining intensity comparisons were performed only between samples within the same set after blinding.

### Image acquisition and analysis

All histological images were acquired using laser-scanning confocal microscopes (Carl Zeiss LSM 710 or LSM 900) with wavelength-appropriate excitation and emission filters, a pinhole setting of 1 Airy unit (AU), and either a 40× oil-immersion objective (Fig5-6) or a 20× objective (Fig 7). Laser power and detector gain were manually adjusted to prevent signal saturation while ensuring clear visualisation of neuronal processes. Images were acquired at 1× zoom (0.415 µm/pixel) with a resolution of 512 × 512 pixels and collected as z-stacks with 1 µm step intervals. Laser power, digital gain, and offset parameters were kept constant for all images acquired within the same set. Image analyses were performed using Fiji (ImageJ; Schindelin et al., 2012). Cells were selected based on TH or tdTomato fluorescence, and the soma outline was manually traced using the freehand selection tool on the single optical section in which the soma displayed the largest cross-sectional area. Mean fluorescence intensity (grey values) for TH and cFos was measured for each cell using the “Measure” function. The mean intensity value for each cell was normalised to the average staining intensity of all measured cells within the control slice of the corresponding co-embedded set (Galliano et al., 2021).

### Statistical analysis

Statistical analyses were performed using R. The biological replicate was defined as the mouse (N), while individual neurons were treated as measurements (n). For comparisons between experimental groups (Sham vs Occluded) in Fig 5, 6 were performed using a linear mixed-effects model using the lmer function in the lme4 package in R with treatment as the fixed effect and a by-mouse as the random intercept to account for variability between mice, with neuronal measurements nested within the corresponding mouse (Aarts et al., 2014). Models were fit by restricted maximum likelihood (REML). Fixed-effect significance was evaluated using t-tests with Satterthwaite’s approximation for degrees of freedom, implemented via the lmerTest package. A Type III analysis of variance with Satterthwaite’s method was used to assess the overall effect of Condition. In Fig 5C, E, to assess the effects of Location and Treatment on fluorescence intensity, a two-way analysis of variance (ANOVA) with Location, Treatment, and their interaction as factors (intensity∼ Location* Treatment) were performed.

In Fig 7, to assess the effect of conditions on fluorescence intensity, a generalised linear mixed-effects model with a Gaussian family and identity link using the glmmTMB package was fitted to summarised observations. For 1-day occluded data, 72 observations, summarised from 6768 TH cells and 6619 cFos cells, across conditions, nested by Image, Slice, Mouse, Set. For 7-day occluded data, 33 observations, summarised from 4507 TH cells, across conditions, nested by Image, Slice, Mouse, Set. Given the hierarchical structure of the data, Image->Slice->Subject->Set, the model included condition as the fixed effect and a nested random-intercept (intensity ∼ Condition + (1|Set) + (1|Set:Subject) + (1|Set:Subject:Slice)). Additionally, to account for heteroscedasticity across images, we included a dispersion model to allow residual variance to differ by Image (dispersion ∼ Image). Fixed-effect significance was assessed using Wald z-tests and corrected for multiple comparisons using two-stage Benjamini-Krieger-Yekutieli method using the mutoss package in R.

Statistical significance was set at α = 0.05, and only significant differences are indicated in figures using asterisks.

## AUTHOR CONTRIBUTIONS

EG designed experiments and supervised the project; AD-R, SP-K, LH, TCS performed experiments; AD-R, LH and SP-K analysed data. AD-R and EG wrote the paper, with all authors contributing to the editing of the manuscript.

## DATA AVAILABILITY

All data, raw imaging data, and R analysis scripts supporting the findings reported in this study will be made publicly available upon acceptance of this manuscript following peer review.

## ACKNOWLEDGEMENTS

This work was supported by a UKRI Medical Research Council Equipment Grant (MC_PC_MR/X012271/1) and project grants from the UKRI Biotechnology and Biological Sciences Research Council (BB/W014688/1) and the Wellcome Trust (301427/Z/23/Z) (EG); a Wellcome Trust Early-Career Award (227621/Z/23/Z; AD-R); a Cambridge Trust PhD Studentship (LH); and a Gates Cambridge Scholarship (TS). We thank Haoming You for technical assistance during plug preparation and Edina Hovarth-Gulacsi for help with mouse colony management and tissue preparation.

